# Constructing spatial perception through self-touch

**DOI:** 10.1101/2020.11.21.392563

**Authors:** A. Cataldo, L. Dupin, H. Dempsey-Jones, H. Gomi, P. Haggard

## Abstract

Classical accounts of spatial perception are based either on the topological layout of sensory receptors, or on implicit spatial information provided by motor commands. In everyday self-touch, as when stroking the left arm with the right hand, these elements are inextricably linked, meaning that tactile and motor contributions to spatial perception cannot readily be disentangled. Here, we developed a robot-mediated form of self-touch in order to decouple the spatial extent of active or passive movements from their tactile consequences. Participants judged the spatial extent of either the movement of the right hand, or of the resulting tactile stimulation to their left forearm. Across five experiments, we found bidirectional interference between motor and tactile information. Crucially, both directions of interference were stronger during active than passive movements. Thus, voluntary motor commands produced stronger integration of multiple signals relevant to spatial perception.

## Introduction

Successful interactions with the environment depend on accurate spatial representations of both the external world and of our body acting upon it. The coding of space is, therefore, an essential requirement for both motor and perceptual systems. Reaching actions, for example, rely on classifying spatial locations into near (e.g., reachable) vs. far space (*1, 2*), and on computing movement vectors to bring the hand to the target location. Similarly, localising tactile stimuli impinging on the skin requires body-centered reference frames based on accurate representations of both the skin relative to the underlying body, and the positions of body parts in space (*3*).

Self-touch is arguably one of the earliest and most ubiquitous spatial experiences. Fetal hand-to-face movements occur in utero from 13 weeks, and the uterine environment makes for frequent interlimb contact (*4*). After birth, several forms of self-touch behaviours persist through childhood and into adulthood, including incidental contact between the hands during bimanual object handling (e.g., tying shoelaces), grooming actions, and functional self-stimulation (e.g., clapping hands to express approval, or to keep warm; grasping a wounded body part). These self-touch behaviours all involve a distinctive sensorimotor contingency between the neural information relating to the moving body part and the stimulation sensed by skin receptors in the touched body part. This situation, often termed *touchant-touché* (i.e., motor *touching* and sensory touch*ed*, respectively) (*5–7*), results in highly correlated motor and sensory representations in the brain, which has been linked to the development of self-awareness (*6, 8*) and body representation (*7, 9–12*) For our purposes, the causal dependence between the motor (*touchant*) and somatosensory (*touché*) components of self-touch means that spatial features are coded twice, by distinct but related spatial codes for action and for perception respectively. For example, if we slide our right index finger along our left forearm, the movement we perform with the finger and the touch we feel on the arm both carry spatial information. Further, this tight spatial relationship between movement and touch contributes to their perceptual fusion into a single psychological event, so that movement and touch are perceived as having the same spatial extent. The double sensation of *touchant-touché* indeed forms a central element in phenomenological accounts of the bodily self (*6*). However, few experimental studies have investigated the relative contributions of motor and sensory information to this integrated percept.

Interestingly, self-touch is widely discussed in early discussions of spatial perception. The origins of our “amodal and invariant sense of space” (*13*) are still a matter of debate. Many neuroscientific discussions of spatial perception stress the orderly topographic projections from receptor surfaces, such as the retina and skin, to the brain (*14, 15*). However, if these projections are taken as explanations of spatial perception, they may seem circular, since they apparently explain (external) space in terms of (internal, neurotopographic) space. In contrast, *local sign* theories of space perception instead explained the sensory quality that a stimulus has in virtue of where it is located (i.e., its “thereness”) in terms of the motor commands required to orient to the stimulus. Thus, perception of visual location was explained in terms of the saccadic motor command required to fixate that location (*16*); while perception of tactile location was explained in terms of the reaching movement required to touch that location (*17, 18*). On this view, active self-touch provides an underpinning mechanism of space perception. However, these descriptions were ultimately thought experiments, and were not accompanied by extensive experimental evidence (*17, 18*).

In our example scenario of stroking the forearm with the finger, local sign theories clearly predict that the spatial nature of the resulting percept comes from the motor command to move the finger, and not from tactile, sensory signals from the skin region that is stroked. Early hypotheses of motor dominance over sensory signals have continued to be influential in psychology and neuroscience of perception, finding modern echoes in theories such as perceptual enactivism (*19*), and active vision (*20, 21*).

Table 1 outlines some alternative theoretical accounts of how spatial percepts might arise in self-touch. First, the brain might maintain completely independent spatial representations for *touchant* (movement) and for *touché* (tactile sensation) (*22, 23*), implying no interference between signals Table 1, Hypothesis A. Second, the motor signal might dominate the tactile signal as suggested by local sign theories, or *vice versa*, producing asymmetric interference between movement and touch in spatial perception (*17, 18*), Table 1, Hypothesis B. Third, motor and tactile signals might fuse, either along the lines of optimal multisensory integration (*24*), or suboptimally, to produce a single spatial percept, Table 1, Hypothesis C. Each theory makes distinct predictions about the weighting that the motor signal will have when participants are asked to report the spatial perception of the touch, and vice ve*rsa*. We will refer to this weighting measure of interference or automatic integration as ω.

**Table 1.**
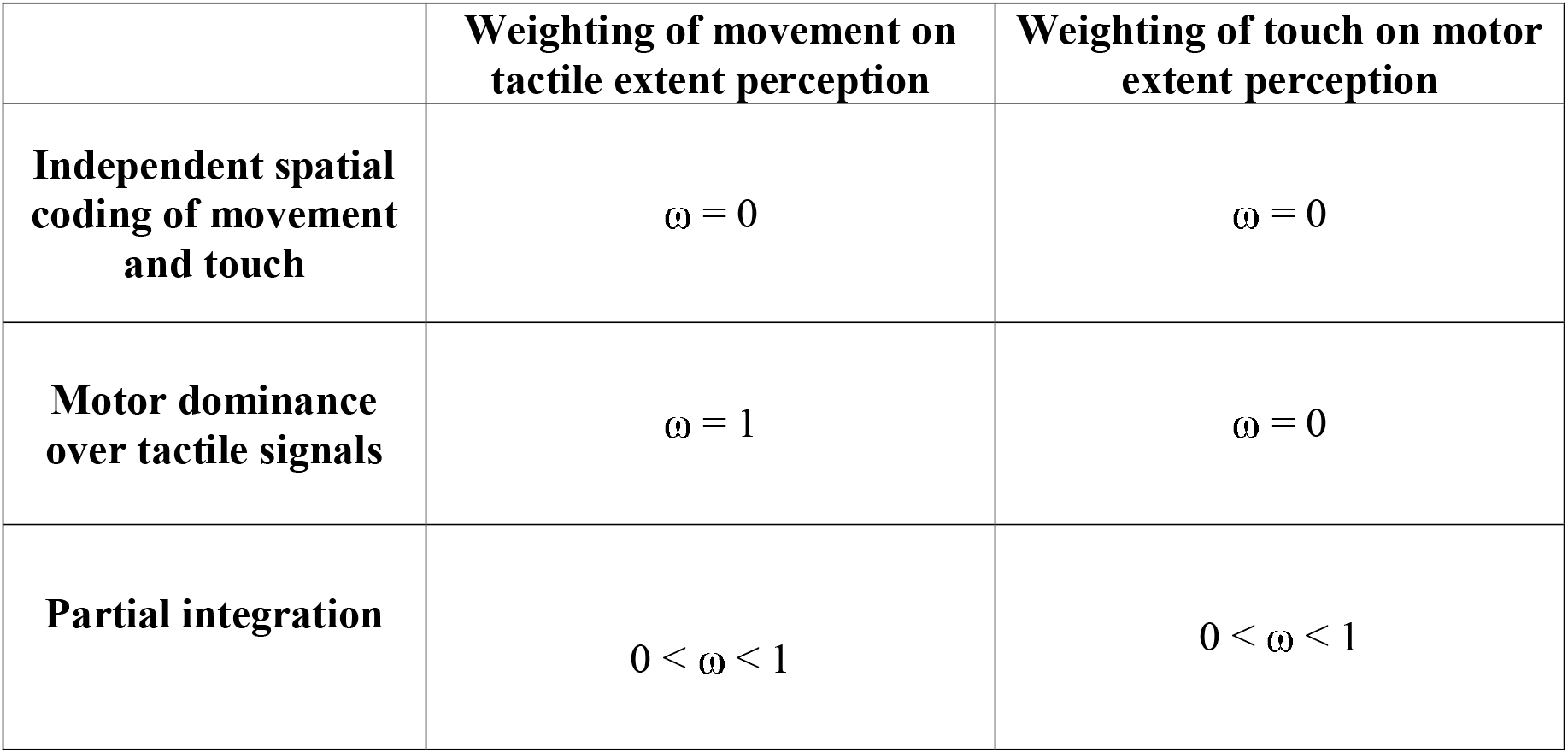
Three alternative accounts of the relation between motor and tactile signals during self-touch, and their predictions for the interference weighting between signals.

Here, we used two robots linked in a leader-follower configuration with a computer-controlled gain between them, to achieve a laboratory version of the common self-touch experience of stroking the left forearm with the right hand. Participants moved the handle of the leader robot with their right hand and simultaneously felt a corresponding stroke on the left forearm from a brush attached to the follower robot (see Figure 1). The computer-controlled gain of the robot coupling allowed the tactile stimulation to be shorter, equal, or longer in extent than the movement that caused it, thus decoupling the normally fixed spatial relation between *touchant* and *touché*. We investigated the patterns of interference between these signals by asking participants to judge either the extent of the tactile stroke they felt (Experiment 1), or the extent of the movement they made (Experiment 2). Participants either actively moved their right hand, or it was passively moved. In the active self-touch condition, participants actively moved the leader robot with their right hand to induce self-touch on their other arm. In the passive self-touch conditions, instead, the participant held the robot with their right hand, while the experimenter passively moved it, again causing a matching self-touch stimulation of the participant’s left forearm. Comparing our active and passive conditions allowed us to investigate the importance of voluntary action to spatial perception. Experiment 3 used a within-subject design, in which the same participants gave both movement judgements and touch judgements. This allowed a stronger within-participant test of asymmetric interference between movement and touch, an opportunity to correlate weights for movement judgements with those for touch judgements, and a direct test of optimal integration theories by relating each signal’s weight to its precision. Finally, two control experiments looked at whether our results could be caused by effects on spatial extent judgement of the different velocities of movement and touch (the ‘tau effect’) (*25*) produced by changing the motor:tactile gain. To do this, we tested unimodal versions of each task, where participants either judged the extent of the tactile sensation without any concomitant movement (i.e. “touch only” condition, Experiment 4), or judged the extent of movement in absence of any tactile stimulation (i.e. “movement only” condition, Experiment 5). As both the control experiments involved unimodal judgements, interference from another signal could not have affected the results.

**Figure 1.**
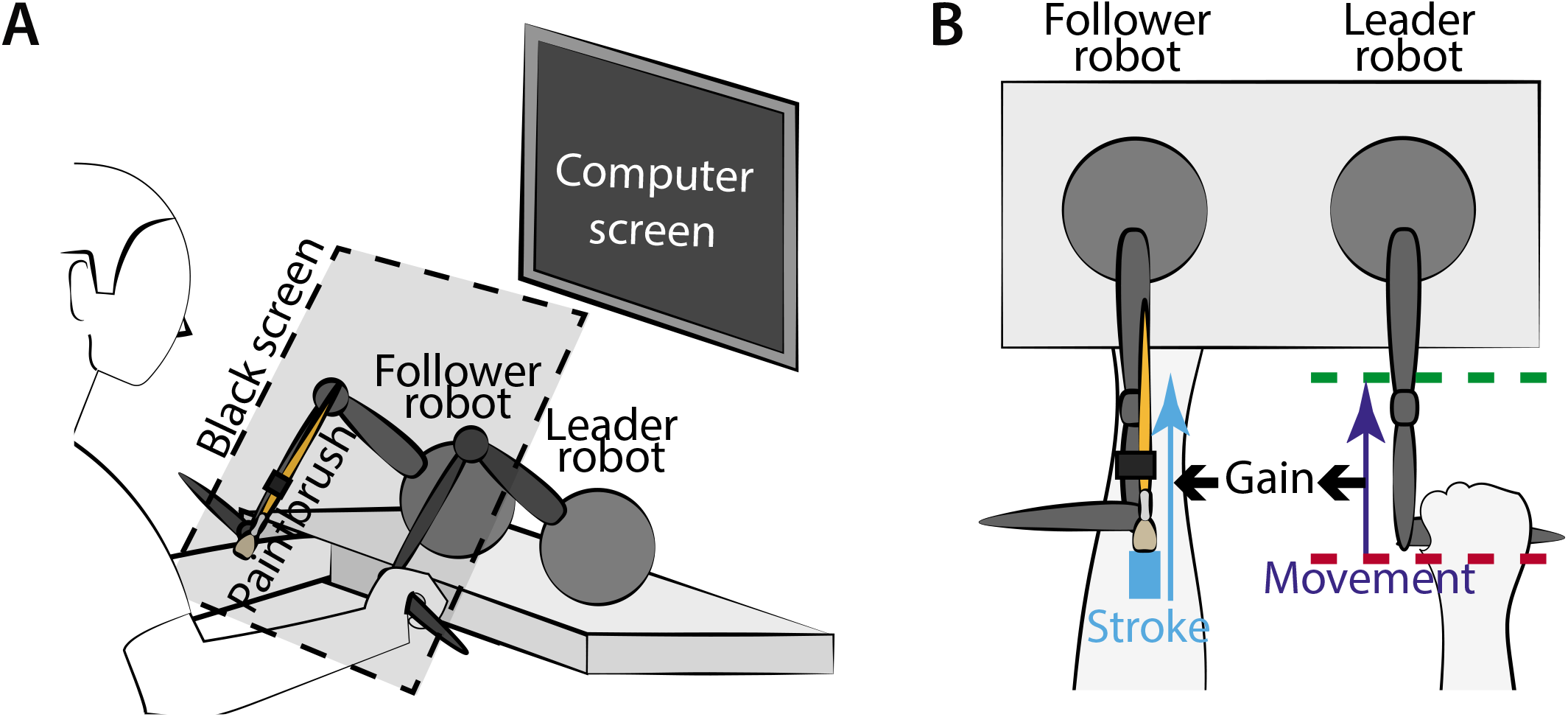
Experimental setup and stimuli. **A.** Participants moved the handle of the leader robot with their right hand and simultaneously felt a corresponding stroke on the left forearm from a brush attached to the follower robot. A black screen (black dashed line) covered both the participants’ arms and the robotic setup throughout the experiment. **B.** The physical extent of right arm movement was modulated via two “virtual walls” defining start (red dashed line) and stop (green dashed line) positions, which varied between trials. The relation between the extent of movement (dark blue arrow) and touch (light blue arrow) depended on the gain of the leader:follower robot coupling, which was randomized across trials (see https://tinyurl.com/yxf34yna for a video of the setup).

## Results

### Overall performance

Perceptual performance was generally good in all experiments. We found a monotonic relation between perceived and actual spatial extent for both movement and touch judgement conditions in both active and passive self-touch (see panels A and B in Figures 2–4 and Supplementary ANOVA Tables S1-5). Participants were thus able to perceive spatial extent in all conditions.

**Figure 2.**
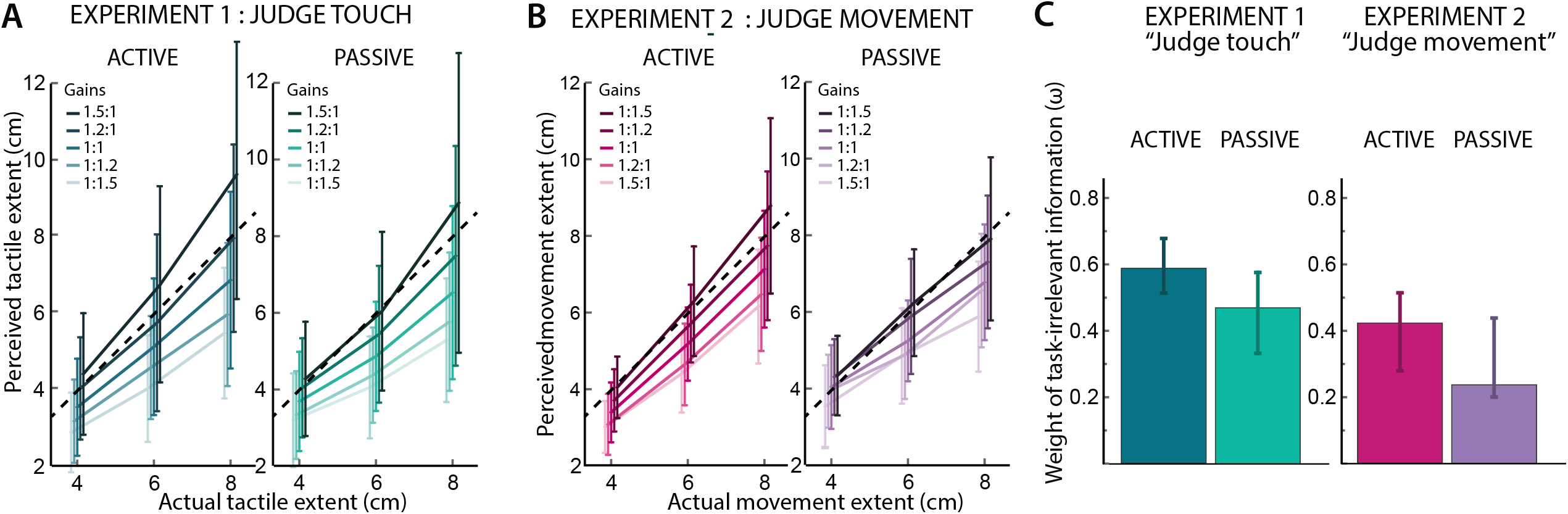
Results from Experiment 1 and 2. **A** Mean perceived tactile extent (Experiment 1) as a function of actual stimulus extent, and gain applied to the task-irrelevant information. **B** Mean perceived movement extent (Experiment 2) as a function of actual movement extent and gain applied the task irrelevant information. Error bars in **A-B** represent the Standard Deviation of the Mean (SD). **C** Median weights (ω) of the task-irrelevant information (median was used because weights were not normally distributed). The positive weights in both experiments show that motor information influences tactile judgement even when task-irrelevant, and that tactile information similarly influences judgements about movement. Error bars represent the 95% CIs for the median (*26*).

**Figure 3.**
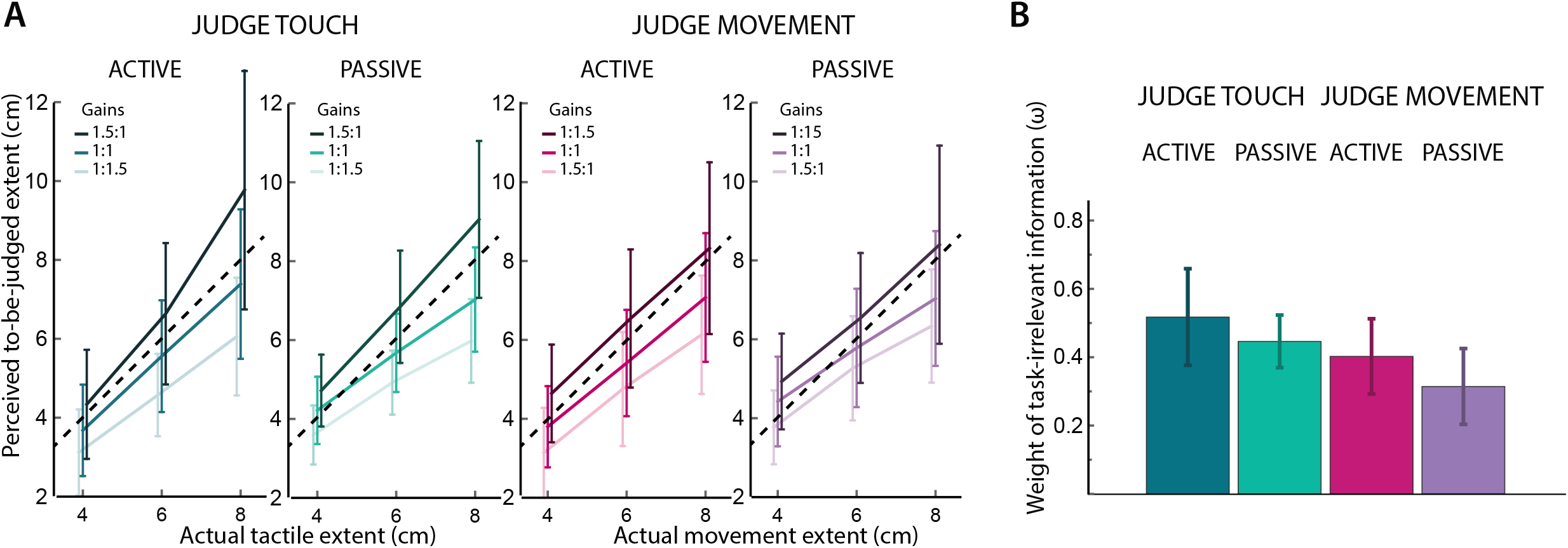
Results from Experiment 3. **A-B** Mean perceived extent of the target sensation as a function of actual stimulus extent, and gain applied to the task-irrelevant information in Experiment 3. Error bars represents the SD. **C** Mean weights (ω) of the task-irrelevant information in Experiment 3. Error bars represent the 95%CI of the mean.

**Figure 4.**
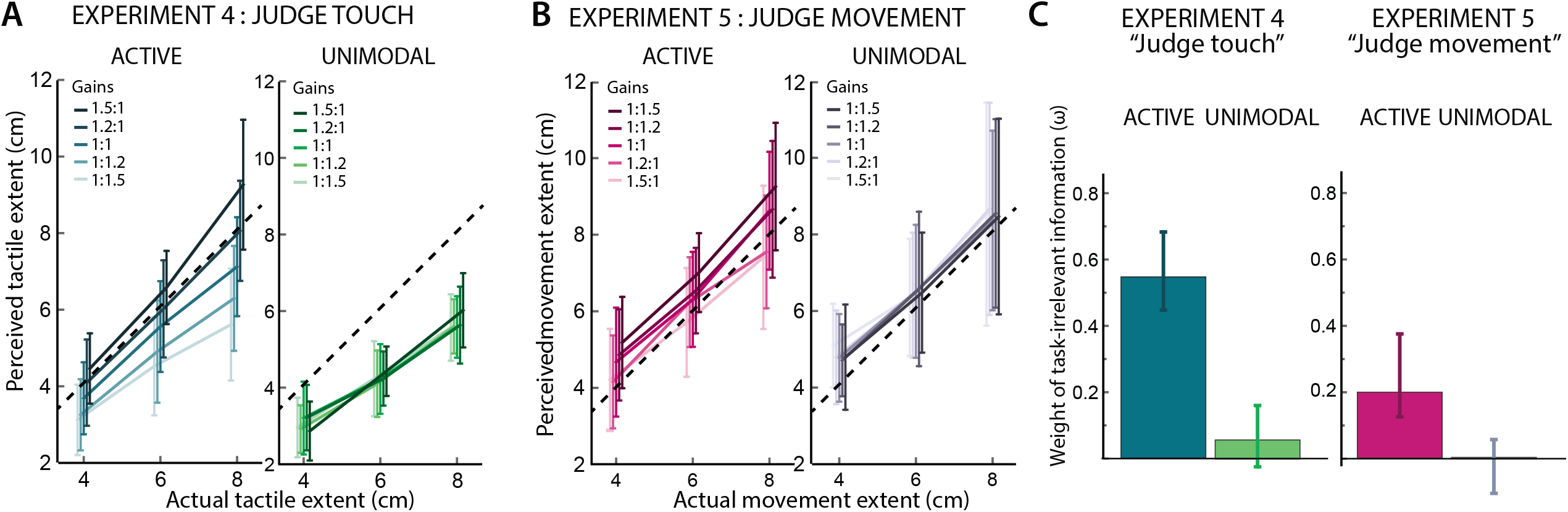
Results from control Experiment 4 and 5. **A-B** In Experiments 4 and 5, the passive conditions were replaced with unimodal versions of the task where tactile (**A**) or movement (**B**) sensations occurred in absence of task-irrelevant information. Error bars in **A-B** represents SD. **C** Weights (ω) of the task-irrelevant information. Error bars represent 95% CIs of the median.

Experimentally manipulating the motor:tactile gain allowed us to quantify the effect of movement information on tactile perception and *vice versa*. Stronger effects of gain manipulations correspond to stronger interference from the task-irrelevant signal on the to-be-judged signal. We therefore used equation 1 (see Statistical Analysis) to compute the weight of task-irrelevant information (i.e. weighting of movement extent when the task was to judge tactile extent and vice versa) in each experimental condition (judge touch/movement x active/passive) and in the unimodal control conditions. The resulting weights (denoted by ω) for each participant and each experiment are given in the Supplementary Materials.

### Are touchant and touché independent?

An account of independent spatial perception for action and tactile perception (*22, 23*) predicts no influence of movement on judgements of tactile extent, and no influence of tactile extent on judgements of movement extent, as the motor:tactile gain is varied (see Table 1A). Our measure of the weight of the task-irrelevant sensation thus provides a summary measure of the effects of changing motor:tactile gain.

Figure 2 A-B shows the mean perceived extents for each to-be-judged information as a function of the actual extent and the gain applied to the task-irrelevant information in Experiment 1 (i.e. “judge touch”; Figure 2A) and Experiment 2 (i.e. “judge movement”; Figure 2B). The data were not normally distributed (see Supplementary Table S11), and were therefore tested using the Wilcoxon’s Sign Test (see Statistical Analysis section). We compared each condition against 0 (where a ω of 0 would indicate no effect of the task-irrelevant information, see Table 1, hypothesis A) and against 1 (where a ω of 1 would indicate complete dominance of one signal over the other, e.g., Table 1, hypothesis B) within each experiment, so we Bonferroni corrected for four comparisons per experiment, giving α = 0.0125 per test.

The weights (ω) of the task-irrelevant information (Figure 2C) were significantly greater that than 0 in both type of task and type of movement (Experiment 1: Judge Touch – Active: median ω = 0.59 [95% Confidence Interval of the median = 0.52, 0.68]; Judge Touch – Passive: 0.47 [0.34, 0.58]; Experiment 2: Judge Movement – Active: 0.42 [0.28, 0.52]; Judge Movement – Passive: 0.24 [0.21, 0.44]; test against 0, *Z* < −3.059, *p* < 0.002, *r* < 0.883, in all cases).

That is, when participants were instructed to judge the spatial extent of the tactile stroking, they were nonetheless influenced by the extent of the movement, and *vice versa*. Thus, the two components of self-touch strongly influenced each other, even when task irrelevant. This finding clearly rejects a hypothesis of complete independence between motor and somatosensory signals in extent perception (Table 1A).

### Do motor signals dominate tactile perception in self-touch?

Motor-based theories of space perception hold that motor signals dominate over sensory signals (*17, 18*). Under this hypothesis, tactile stroking should thus have little to nil influence on perception of movement (i.e., a weight *ω* = 0 for touch in the “judge movement” task, see Table 1A). Conversely, movement should strongly influence, or even totally dominate tactile extent perception (i.e., a weight *ω* = 1 for task-irrelevant movement in the “judge touch” task, Table 1B). Our previous analysis provides evidence against the first prediction of motor dominance theories, by showing that the weights of tactile information were significantly higher than 0. Similarly, contrary to the second hypothesis of the motor dominance theories, the effect of movement on touch was significantly different from a prediction of total dominance, since all ωs were significantly lower than 1 (*Z* < −2.824, *p* < 0.005, *r* < 0.88, in all cases; Bonferroni adjusted for four multiple comparisons: α = 0.0125 per test).

Thus, our results suggest that theories that reduce spatial perception to motor signals cannot readily account for the bidirectional interference in spatial extent perception during our self-touch manipulation.

### Partial integration of motor and tactile information during self-touch

The results of Experiments 1 and 2 suggest that both tactile and motor information are partially integrated during self-touch. To investigate this partial integration further, and to directly compare the weights of the irrelevant information in the two tasks (“judge touch”, “judge movement”) we asked a new group of participants in Experiment 3 to judge both movement and touch extents, in separate blocks. In Experiments 1 and 2, direct comparison of weights would include an element of inter-participant variability, because of the between-subjects comparison. Only three motor:tactile gains were tested, but these spanned the same range as Experiments 1 and 2 (see Methods). Mean perceived to-be-judged extents for each gain are presented in Figure 3A.

First, we analysed Experiment 3 to replicate the results of Experiments 1 and 2. As the data were normally distributed (see Supplementary Table S6), we used t-tests to analyse the weights. As in the previous experiments, all weights *ω* were significantly greater than 0 (t_23_ > 5. 99, *p* < 0.001, Cohen’s *d* > 1.22, in all cases) and lower than 1 (t23 < −10.001, *p* < 0.001, Cohen’s *d* > 2.042, in all cases, Bonferroni adjusted for two comparisons in each of four conditions, i.e., α = 0.0063 per test) (see Figure 3B) (Judge Touch – Active: mean ω = 0.52 [±95% CI = 0.09]; Judge Touch – Passive: 0.45 [0.07, 0.17]. Judge Movement – Active: 0.4 [0.1, 0.24]; Judge Movement – Passive: 0.31 [0.11, 0.28]).

Next, to test for effects of type of judgement (movement extent, tactile extent) and type of movement (active, passive), and directly compare weights across these conditions, we used a 2 x 2 repeated measures ANOVA. The ANOVA showed a significant main effect of type of Task (F_1_,_23_ = 5.21, *p* = 0.032, η_p_^2^ = 0.19) with a greater weight of movement when participants had to judge touch (mean *ω* ± 95% CI: 0.48 ± 0.08) than *vice versa* (0.36 ± 0.10), indicating a directional asymmetry in the interference between movement and touch signals. Moreover, there was also a 2 significant main effect of Movement type (F_1,23_ = 10.1, *p* = 0.004, η_p_^2^ = 0.31) with higher weights, indicating stronger interference, when movement was active (mean *ω* ± 95% CI: 0.46 ± 0.08) than passive (0.38 ± 0.07). There was no significant interaction between the two factors (F_1,23_ = 0.17, *p* = 0.68).

Finally, no correlation between the interference of movement on touch and of touch on movement was found, in either active or passive conditions (see Supplementary Figure S1 and explanatory text in Supplementary Material). However, Active and Passive movement conditions in both tasks were strongly correlated (see Supplementary Figure S2 and explanatory text in Supplementary Material), suggesting that, within each task, the influence of irrelevant information occurs due to some process that is common to active and passive conditions.

Thus, these results show strong and bidirectional, but asymmetrical interference between the *touchant* and the *touché* sensations in self-touch. The interference of movement on tactile extent judgements was greater than the interference of touch on movement extent judgements.

### Self-touch as optimal integration

Current theories of multisensory perception focus on optimal integration of multiple sources of information (*24*). Each source is weighted according to its reliability or precision (*27–29*). To test whether the weightings of tactile and movement signals in self-touch are optimally integrated, we calculated participants’ *precision* for each condition of each experiment (see Statistical Analysis section and Table 2). If optimal integration of tactile and motor information takes place in self-touch, then our weighting values should directly follow from precision data, with precision data showing the same significant main effects of task and movement as weightings, and no interaction.

**Table 2.** Precision data (cm^-2^) for each condition of each experiment.

| Experiment | Task | Condition |  |  |
| --- | --- | --- | --- | --- |
| | | Active movement<br>mean ± SD | Passive movement<br>mean ± SD | Unimodal<br>mean ± SD |
| 1 | Judge Touch | 1.78 ± 1.19 | 1.77 ± 0.90 | |
| 2 | Judge Movement | 1.53 ± 0.49 | 1.39 ± 0.64 | |
| 3 | Judge Touch | 1.37 ± 1.01 | 1.33 ± 0.84 | |
| | Judge Movement | 1.21 ± 0.97 | 1.16 ± 0.57 | |
| 4 | Judge Touch | 1.8 ± 0.94 | | 2.14 ± 1.40 |
| 5 | Judge Movement | 0.94 ± 0.48 | | 1.08 ± 0.61 |

Yet, a 2 x 2 repeated measures ANOVA on the precision data of Experiment 3 showed no significant main effects (type of Task: F_1,23_ = 1.37, *p* = 0.25; type of Movement: F_1,23_ = 0.15, *p* = 0.71) nor interaction (F_1,23_ = 0.002, *p* = 0.96). Similarly, we did not find any difference in precision between the type of movement in Experiment 1 and 2 (t_11_= 0.06, p = 0.96 and t_11_= 0.92, p = 0.38 respectively using paired t-test), nor between active and unimodal conditions in Experiments 4 and 5 (t_11_= 1.05, p = 0.32 and t_11_= 0.81, p = 0.41 respectively using paired t-test). Thus, the weightings for interference between movement and touch do not simply follow the precision of the component signals.

### Could the “tau effect” explain our results? Control experiments

Since the movement durations of the leader and follower robots were matched, changes in motor:tactile gain necessarily modify velocity of the tactile stroking stimulus. Extent perception can be affected by the velocity or duration of the stimulation, a phenomenon often referred to as “tau effect” (*25*). Because in our design movement and touch began and ended together, a high motor:tactile gain would result in both a greater tactile spatial extent *and* a higher tactile velocity, than a low motor:tactile gain. We therefore investigated whether the effects of gain on spatial extent perception truly reflected interference *between* the two different signals, rather than influence of velocity on extent perception *within* a single sensory channel. Therefore, we ran two additional control experiments to investigate whether a tau effect could explain the results of Experiment 1-3.

The active movement conditions of Experiments 4 and 5 were identical to the active conditions of Experiments 1 and 2, and to the corresponding conditions of Experiments 1-3. In a second, “unimodal”, condition, participants were asked to judge the tactile extent in absence of any movement (Control Experiment 4) or, to judge the extent of the movement in absence of any tactile stimulation (Control Experiment 5). In these conditions, any spatial perception is both unimodal and unimanual, since only touch (Experiment 4) or only movement (Experiment 5) is present, and thus there is no interference between movement and touch. If the apparent influence of irrelevant signals in Experiments 1-3 was in fact due to an artefact of the tau effect, then the tau effect should be equally present in unimodal conditions, and our *ω* measure should again be greater than 0. Conversely, if the interference in Experiments 1-3 indeed represents interference from irrelevant information, rather than variations in the velocity of the judged signal, then *ω* in unimodal conditions should show not be different from 0. Participants in Control Experiments 4 and 5 judged spatial extents of touch, and of movement respectively, in two conditions: a unimodal condition described above, and an active self-touch condition replicating Experiments 1-3 (Figure 4A, B).

Data violated the normality assumption (see Supplementary Table S6), and were therefore tested using the Wilcoxon’s Sign Test. In the “unimodal” conditions of both experiments 4 and 5, weights were not significantly different from 0 (Judge Touch – Unimodal: 0.06 [-0.03, 0.15], *Z* = −1.80, *p* = 0.071; Judge Movement – Unimodal: 0.0005 [-0.11, 0.05], *Z* = 0.078, *p* = 0.94; see Figure 4C). Thus, we found no significant evidence for a tau effect, and no reason to attribute the interference effects of Experiments 1-3 to this source. The results for the active condition replicated the effects of Experiment 1-3 (see Supplementary Results).

## Discussion

Self-touch is widely thought to be an important psychophysiological event, but it has rarely been studied experimentally. While other studies have examined the consequences of self-touch (30, 31) and the processing of self-generated stimuli (32), our results provide a systematic experimental manipulation of self-touch stimulation, and a novel focus on spatial perception of self-touch events themselves.

We used an innovative method that allows variable coupling between two haptic robots. We could thus break the direct relation, between hand movement and tactile stimulation of another body part, that characterizes normal self-touch. Participants made voluntary movements of their unseen right hand or were passively moved through an equivalent trajectory. Crucially, they could not predict or decide in advance the amplitude of these movements, which instead depended on haptic walls generated by the robot interface. The movements directly caused a simultaneous, unseen stroking stimulus along the left forearm, in the same direction as the movement, via a leader:follower robot arrangement. The gain of the robotic coupling varied unpredictably across trials, so that the spatial extent of movement and the spatial extent of touch could be decoupled, in contrast to their strict correspondence during natural self-touch. When participants were asked to report the spatial extent of their movement, their perceptual judgements were strongly influenced by the tactile stimulus extent, and vice versa. This automatic perceptual interference, in both directions, from an irrelevant signal was stronger when participants made active movements then when they made passive movements. Control experiments confirmed that these results reflected interference between representations of extent, rather than a confound between extent and velocity introduced by our manipulations of motor:tactile gain and leading to a tau effect. Overall, our results provide robust experimental support for a degree of mutual interference between *touchant* and *touché* during self-touch. In experiments 1-3 both *touchant* and *touché* signals were always present, and always simultaneous, yet participants judged just one of these signals. Our task thus required selective attention. The interference from the unattended signal can be considered either as a limitation of selective attention, or as an automatic, pre-attentive integration between the two signals. However, other studies reported minimal bilateral interference (*33*), and even enhancement (*34*), in many somatosensory perceptual tasks (see *35* for a review), making it unlikely that our results reflect general inability to direct attention. Moreover, general effects of attention cannot readily explain the consistent and strong differences in *touchant*-*touché* interference that we observed for active *vs* passive movement – a point to which we will return below.

Several neurocognitive theories make contrasting predictions about perceptual experience in *touchant-touché* scenarios. First, local sign theories (*17, 18*) perhaps have the most direct relation to our self-touch scenario, because they specifically posit a motoric basis for spatial percepts. These theories specifically predict a strong influence of movement extent on tactile perception, without any reverse influence of touch on movement (*17, 18*). In a strong version of this theory, the motor command signal simply *is* the basis of spatial experience. The weightings of Figure 2C should then be 1 for effects of active movement on touch, and 0 in all other conditions. Indeed, many theories emphasise the dominant role of active motor signals in perception (*24*). Our results do not support an account of total motor dominance, for two reasons. First, the effects of touch on movement perception were clearly significant – event though they were less than the effects of movement on touch perception. This occurred both when a voluntary motor command was present, and, in our passive condition, when it was not. Thus, while movement did influence touch more than touch influenced movement, as predicted by local sign theory, the exclusive reduction of spatial perception to non-spatial, intensive (i.e. based on intensity), as opposed to “extensive” (or based on extent), motor command signals, was not confirmed.

Second, active motor commands lead to an increased bidirectional interference, from touch to movement, as well as from movement to touch, relative to passive movements. A key result of our study, replicated across Experiments 1-3, is thus an increased integration between movement and tactile signals during active vs passive movement, rather than a simple enhancement of motor dominance.

Self-touch is often cited in support of theories of motor-based space (*36, 37*). Further, self-touch is considered a basis of bodily self-awareness and wider spatial cognition (*6, 8*), i.e., that one’s own body is a volumetric spatial object within an external world consisting of other objects. Our results cast doubt on the idea that motor commands form the underpinning foundation of the experiences of tactile space, or of bodily self-awareness, since tactile sensations had a strong reciprocal influence on awareness of movement. Of course, we cannot exclude the possibility that tactile spatial perception initially depends on movement, but later becomes an independent experience through repeated motor-tactile association (*17, 18*). However, such associationist theories would presumably predict that the primary original signal (motor) should continue to dominate the secondary, associated signal (tactile) when both are present – yet we found robust effects of tactile stimulation on perceived movement extent.

Could the array of tactile receptors in the skin then be alone sufficient for spatial perception? This point is controversial: some theories deny the existence of a ‘tactile field’ analogous to the visual field, and hold that the spatial properties of tactile sensation do not reflect signals from tactile receptors themselves, but rather depend on movements that generate specific patterns of tactile contact with external objects (*38*). Conversely, we have previously shown that passive touch on an immobile skin surface is sufficient to develop a rich spatial percept (*39*). Specifically, passive touch supported the same processes of path integration and shape representation that are conventionally used to identify cognitive maps of external space in the navigation literature (*39*).

Other neurocognitive theories also make predictions about self-touch. Neurocomputational models of predictive motor control (*32*) suggest that perceptual consequences of self-generated motor actions are suppressed, or at least attenuated, by being cancelled against predictions of an internal model. On a strict version of this view, the *touché* part of self-touch should generate no sensation at all – yet everyday experience shows this is not the case. In our experiments, however, random trial-to-trial variation in motor:tactile gain meant that tactile stimulation was not entirely predictable from the motor activity. Nevertheless, motor prediction theories would struggle to explain our finding that the sensory consequences of self-touch strongly influenced the perception of voluntary movement extent.

Ideomotor theories of action (*40*) seem to make the opposite prediction, suggesting that actions are mentally represented in terms of their external outcomes, rather than the internal motor commands used to achieve those outcomes. On this view, one might expect tactile signals to dominate self-touch perception, yet this was not found.

Finally, multisensory integration theories suggest that signals relating to a common source are integrated to provide a single representation. Optimal integration theories predict that more reliable (precise) signals should be more strongly weighted. Previous observations of strong integration and coherence during *touchant-touché* sensations (*6*) suggests these models might apply to self-touch. Our methods used automatic interference of an irrelevant signal on a to-be-judged signal, rather than reports of a common source event traditionally used to study optimal integration (*24*). Nevertheless, the mutual influences on movement on touch perception and *vice versa* that we observed did not appear to follow predictions of optimal integration theory. First, we found that movement information was more highly weighted than tactile information in our main Experiments 1-3. Optimal integration theory would predict this pattern to reflect a higher precision for judgements of movement extent compared to tactile extent, but we observed a (non-significant) difference in the opposite direction. Thus, while we did not formally use multisensory integration framework for our study, the pattern of interference between self-touch signals that we found points to suboptimal integration.

An interesting feature of our results was the greater bidirectional interference between motor and tactile signals in active, compared to passive movement. Local sign theories would predict a stronger influence of active compared to passive movement on judgements of touch, but a weaker influence of touch on judgements of active compared to passive movement. In fact, we found both increased influence of movement on tactile judgement, and also increased influence of touch on movement judgement, for active compared to passive conditions. The latter finding seems in stark contradiction to local sign theory, and models of ‘motor-based space’ generally. While the increased interference during active compared to passive movement was not originally predicted for Experiments 1-2, the effect was strongly replicated using a larger sample size and a within-participant design in Experiment 3. Our results thus suggest that presence of voluntary motor commands led to an increased interaction between movement and tactile signals, implying stronger automatic integration. At first sight, this might simply seem an effect of selective attention to action. When participants must additionally focus on controlling their right hand movement, they might be less able to attend to other signals such as the tactile stimulation of the left hand. Stronger attentional demands of active *vs* passive movement could potentially explain the stronger interference of active *vs* passive movements on judgements of touch. However, those same attentional demands of active movement cannot also explain the stronger interference from touch on judgements of active compared to passive movements. Further, this hypothesis would predict low precision of touch judgement in active vs. passive movement. However, we did not find any significant effects of active versus passive movement on perceptual precision across our five experiments.

Instead, we suggest that the presence of a voluntary motor command may promote integration between multiple sensory and motor signals present in self-touch. A distinctive feature of voluntary movement is its instrumentality: voluntary actions often aim to achieve a specific outcome. Action and outcome are then represented as bound together, as suggested by ideomotor and reinforcement learning theories (*40–42*). Such binding processes imply a readiness-to-associate of voluntary action. By this we mean that voluntary motor commands should readily integrate with signals carrying information about the consequences of the action. Across three experiments, the presence of a voluntary motor command lead to an increased influence of movement on touch, but also to an increased influence of touch on movement. We therefore suggest that voluntary actions have a distinctive psychological effect of promoting multisensory binding between diverse signals, to produce more integrated, coherent representation of action events. Previous studies have suggested similar integrative functions of voluntary action, in bodily illusions (*43*) and in time perception (*44*). Our result adds a novel dimension to this general view of the integrative nature of voluntary action. It may also explain how voluntary action contributes to the experience of one’s body as a coherent, unified self-object, despite the striking diversity of sensory signals reaching the brain from the body.

Our study has several limitations. First, we studied only the *spatial* aspects of self-touch. Our results cannot therefore address other important aspects of self-touch, such as the self-other distinction (*45*). Second, we used a measurement framework based on selective influence and interference between signals, rather than an optimal multisensory integration framework. Therefore, we cannot formally establish whether *touchant* and *touché* signals are integrated in a mathematical sense. An integration framework would imply asking participants to report a *single* percept (e.g., “What was the extent of that self-touch event?”), whereas our primary concern was to establish the different perceptual contents associated with each individual component signal in the *touchant*-*touché* situation. Nevertheless, the results of our study are consistent with a strong integration of these signals, while suggesting that the integration process itself may be suboptimal.

To conclude, we reported several experiments on spatial perception during self-touch. Novel experimental manipulations of the relationship between movement and touch allowed us to investigate the contributions of each signal to spatial perception, and the degree of interference between one signal and another. We found strong interference of movement on judgements of touch, and also of touch on judgements of movement. While motor signals dominated tactile signals in self-touch processing, classical local sign theories and motor-based space theories are not consistent with the strong interfering effects of tactile input on perceived extent of movement that we repeatedly found. Further, interference effects in both directions were enhanced under active voluntary movement, compared to passive movement. This suggests that a distinctive cognitive consequence of the voluntary motor command is to promote a general integration of multiple signals to synthesise representations related to the body, and thus produce a coherent overall representation of the bodily self. In this sense, our results reveal a simple sensorimotor basis for the intuition that voluntary action underlies the coherence and unity of self-awareness. We have proposed that the voluntary motor command induces a cognitive binding process that facilitates perceptual integration of multiple signals. The mechanisms underlying this distinctive feature of volition, and its relation to phenomenal self-models (*46*) remain unclear, but action-outcome learning is likely to provide a key mechanism (*47*).

## Materials and Methods

### Participants

The sample size for experiment 1 (n = 12: 7 females, mean age ± SD: 22.7 ± 3.1) was decided a priori on the basis of previous similar studies (*7, 12*). To determine the sample size for experiments 2-5, we performed a power analysis based on the results of Experiment 1. The effect size for the main effect of the robotic gain manipulation in experiment 1 was η^2^ = 0.722 (see Supplementary ANOVA Table S1 in the Supplementary Material), considered to be very large using Cohen’s criteria (*48*). With an alpha = 0.05 and power = 0.8, the projected sample size indicated to demonstrate interference effects on movement on touch perception and *vice versa* was 4 participants (G*Power 3.1.9.2 software) (*49*). We nevertheless set a sample size of n = 12 for Experiments 2, 4, and 5 (Experiment 2: 11 females, mean age ± SD: 25.2; Experiment 4: 8 females, mean age ± SD: 24.4 ± 3.8; Experiment 5: 8 females, mean age ± SD: 24.1 ± 3.6), and of n = 24 for Experiment 3 (17 females, mean age ± SD: 29.5 ± 13.2). Eighty-two healthy right-handed volunteers were originally recruited for the study. Two participants were excluded because of technical issues. Based on a priori exclusion criteria, eight further participants were excluded during the training phase because they proved unable to use the robotic device to produce smooth self-stimulation movements (see procedure below). The experimental protocol was approved by the Research Ethics Committee of University College London and adhered to the ethical standards of the Declaration of Helsinki. All participants were naïve regarding the hypotheses underlying the experiment and provided their written informed consent before the beginning of the testing, after receiving written and verbal explanations of the purpose of the study.

### Apparatus

Figure 1 shows a schematic representation of the setup. Participants sat in front of a computer screen with their left arm on a fixed moulded support, and their right arm on an articulated armrest support (Ergorest, series 330 011, Finland). Both the participants’ arms and the robotic setup were covered by a horizontal screen and remained unseen throughout the experiment. The sensorimotor self-touch stimulation was implemented using two six-degrees-of-freedom robotic arms (3D Systems, Geomagic Touch X, South Carolina, USA) linked as a computer-controlled leader-follower system. In this system, any 3D-movement of the right-hand leader robot is reproduced by the follower robot. The estimated lag between the robot trajectories was 2.5 ms (see Supplementary Methods for details). The follower robot carries a paintbrush (12.7 mm width) that strokes the participant’s left forearm (see https://tinyurl.com/yxf34yna for a video of the setup). This setup allowed us to manipulate the gain between the leader and the follower robots so as to produce different combinations of motor and tactile displacements. For instance, if the motor:tactile was set to 1:1.5 then every 1 cm movement of the leader (movement) robot would cause 1.5 cm movement of the follower robot.

The extent of each movement in the anteroposterior direction was controlled by two “virtual walls” created by the force-feedback system of the leader robots. That is, participants would move the leader arm forward/backward until resistance from the force feedback wall prevented them from moving further. This allowed the movement extent to be randomized across trials.

### Experimental design

Experiments 1 and 2 tested, respectively, the effect of movement on tactile extent (judge touch task) and vice versa (judge movement task). Each experiment had a 2 (movement type: active, passive) x 3 (extent of the to-be-judged stimulus: 4, 6, 8 cm) x 5 (motor:tactile gains: 1.5:1, 1.2:1, 1:1, 1:1.2, 1:1.5) within subject design. The movement type factor (active/passive) was blocked and counterbalanced across participants. The spatial extent of the to-be-judged events (movement, or stroke) was randomised. Each of the 30 possible combinations of these factors was experienced eight times, giving a total of 240 trials per participant. The testing was divided into 16 blocks of 15 trials each, and breaks were allowed between blocks.

Experiments 1 and 2 tested for the effects of movement extent on judgements of tactile extent, and for the effects of tactile extent on judgements of movement extent, respectively. Five levels of motor:tactile gain were tested, in randomized order.

Experiment 3 used a full within-subjects design with a 2 (judgement type: judge movement, judge touch) x 2 (movement type: active, passive) x 3 (extent of the to-be-judged stimulus: 4, 6, 8 cm) x 3 (motor:tactile gains: 1.5:1, 1:1, 1:1.5) paradigm. Each of the resulting 36 conditions was repeated eight times, for a total of 288 trials per participant. The testing was divided into 16 blocks of 18 trials each, and breaks were allowed between blocks.

Experiments 4 and 5 aimed to control for the contribution of differential stimulus velocity produced by the gain manipulation and were based on the same experimental design as Experiments 1 and 2. In these experiments, the passive movement condition was replaced with a purely unimodal, and unimanual condition in which participants judged the extent of either touch (Experiment 4) or movement (Experiment 5) in absence of any movement, or of any tactile stimulation respectively.

### Procedure

Participants were familiarised with the experimental setup at the beginning of the experiment, and received training before each condition. In the active movement condition, participants were instructed to perform a back-and-forth movement of the right hand from the far wall to the near wall, and then returning to the starting position (far wall). Participants would move their hand forward/backward until they discovered the position of the virtual walls on each trial, guided by the haptic feel of force when they touched the wall. This was followed by a short auditory beep, as an additional cue they had reached the wall. In the passive movement condition, the handle of the leader robot was moved by the experimenter in the same back-and-forth trajectory described for the active condition. Participants held the leader robot’s handle with their right hand and followed passively the movements produced by the experimenter. The unimodal conditions (Experiments 4-5) were identical to the passive conditions in Experiments 1-2, with the only exception that the participant kept the right, active movement hand (Experiment 4) or the left, passive touch recipient hand (Experiment 5) on the desk, away from the setup. Participants were, thus, judging the extent of touch in absence of movement or vice versa.

Each training phase ended with a practice block of the spatial extent judgement task. Participants were asked to focus only on the “to-be-judged” experience of the block – either the extent of the right hand’s movement, or the extent of the stroke on the left forearm, as appropriate – and to ignore the other sensation. After each active or passive movement, the fixation cross on the screen was replaced by a line of a random length (between 2 and 10 cm). Participants then used two foot-pedals (one which made the line longer, and the other shorter) to adjust the length of the line on the screen. Their task was to match the line on the screen to the extent of either the movement or the tactile sensation, depending on condition. The fixation cross and judgement task line were aligned with the participants’ left arm in the case of the “judge touch” task, and with the participants’ right hand in the case of the “judge movement” task. After adjusting the length of the line, participants clicked a button on the handle of the leader robot to confirm their response and start a new trial.

In all trials of the practice block, movements and tactile feedback were 8 cm in length, so the spatial extent of natural self-touch was consistent with movement extent, as in typical self-touch. The main testing phase was identical to the training phase, except that the gain between the leader and follower robots was systematically manipulated in order to obtain different extents for movement and touch sensations. The gain varied randomly across trials between the different gain values set in each experiment (see above). Thus, although a general consistency between movement and tactile extents remained, participants could not reliably predict movement extent from tactile extent or vice versa. This allowed us to investigate the perceived extent of the to-be-judged sensation (e.g. touch in the “judge touch” task), as a function of the task-irrelevant spatial extent of the other, task-irrelevant event (e.g. movement in the “judge touch” task).

### Statistical analysis

#### Weight of task irrelevant information

The main goal of this study was to test some of the most influential accounts of tactile self-touch space perception using a self-touch paradigm. In particular, we contrasted the theory of independent coding, motor dominance, and partial fusion between motor and tactile spatial information. Each of these groups of theories makes clear predictions (see Table 1) on how the sensory and motor components of self-touch would be weighted according to the task demands (judge touch/judge movement) and the type of movement (active/passive). First, a model based on independent spatial coding for action and perception (*22, 23*) predicts that participants’ extent judgements in each task should be unaffected by the task-irrelevant information. Second, a strong motor-based theory of space perception (*17, 18*) predicts a strong influence of movement on tactile extent (i.e. a weight *ω* > 0 for movement in the “judge touch” task), but much less, or zero, influence of tactile stroking on perception of movement (i.e. a weight *ω* = 0 for touch in the “judge movement” task). Finally, a weight *ω* in between 0 and 1 would suggest partial integration. This could be either optimal (*24*) or sub-optimal, depending on whether the weighting is determined by the precision of each modality or not.

To test these hypotheses, we first used a regression approach to extract a summary measure of sensitivity describing the relation between perceived and actual extent of stimulus. In particular, we fitted the following model to quantify the effect of the task-irrelevant extent information on the to-be-judged extent.

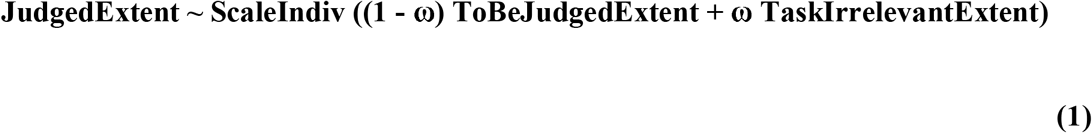

Where ScaleIndiv is an individual scaling factor to capture each participant’s cross-modal mapping from motor/tactile stimulation extent to visual line response, ω is the weight of the task-irrelevant extent (TaskIrrelevantExtent) on the judged extent (ToBeJudgedExtent) information. We did not fit any intercept in this model, since we assumed a judged extent of zero in the absence of any actual spatial stimulation (*50*, for a similar approach in perceptual judgement task, see *51*). In this model, a weight ω = 0 would correspond to the situation where the participant would report the target extent independently from the task-irrelevant information (e.g. no effect of movement on touch in the “judge touch” task). Conversely, ω = 1 would mean that the participants’ to-be-judged extent perception is entirely based on the task-irrelevant information and not at all on the to-be-judged information. Finally, a weight between 0 and 1 would indicate the partial integration of task-irrelevant information in judged extent. Fitting this model allowed us to calculate a single summary numerical value from all the raw judgement data, capturing the influence of movement on touch, and another value capturing the influence of touch and movement (see Supplementary Material for individual weights for each participant in each experiment).

#### Precision

For each participant, and each condition, we computed the precision (Precision = 1/Standard Deviation) for each combination of extent and gain. Since our interest focussed on the conditions, while extent and gain were effects of no interest, we then averaged the precisions values across the different levels of extent and gain, to obtain a single mean precision value for each participant and each condition. The mean precision and its standard deviation over participants are shown in Table 2.

#### Normality of data

Data from Experiments 1-2 and 4-5 violated the normality assumption (see Supplementary Material), therefore the different predictions were tested with a series of Wilcoxon’s Sign tests contrasting the weight of the task-irrelevant information in the four conditions (type of judgement x type of movement) against 0 or 1. Data from Experiment 3 were normally distributed, thus the same analysis were conducted using a series of t-tests contrasting the weight of the task-irrelevant information in the four conditions (type of judgement x type of movement) against 0 or 1, and a 2 x 2 repeated measures ANOVA with factors of Judgement type (“judge touch”, “judge movement”) and Movement type (active, passive).

## Author contributions

AC: Conceptualization, Methodology, Software, Formal Analysis, Investigation, Data Curation, Writing – Original Draft, Visualization. LD: Conceptualization, Methodology, Software, Formal Analysis, Writing – Review & Editing. HDJ: Data Curation, Writing – Review & Editing. HG: Resources, Writing – Review & Editing. PH: Conceptualization, Resources, Writing – Review & Editing, Supervision, Project Administration, Funding Acquisition.

## Data and materials availability

The datasets generated during and/or analyzed during the current study are available on https://tinyurl.com/y3ssgho4. All data needed to evaluate the conclusions in the paper are present in the paper and/or the Supplementary Materials.

## References

1. G. Rizzolatti, L. Fadiga, L. Fogassi, V. Gallese, The space around us. Science. 277, 190–191 (1997).

2. N. P. Holmes, C. Spence, The body schema and multisensory representation (s) of peripersonal space. Cognitive processing. 5, 94–105 (2004).

3. E. Azañón, M. R. Longo, S. Soto-Faraco, P. Haggard, The posterior parietal cortex remaps touch into external space. Current Biology. 20, 1304–1309 (2010).

4. A. Kurjak, G. Azumendi, N. Vecek, S. Kupesic, M. Solak, D. Varga, F. Chervenak, Fetal hand movements and facial expression in normal pregnancy studied by four-dimensional sonography. J Perinat Med. 31, 496–508 (2003).

5. S. Bolanowski, R. Verrillo, F. McGlone, Passive, active and intra-active (self) touch. Behavioural brain research. 148, 41–45 (2004).

6. M. Merleau-Ponty, Phénoménologie de la perception (1945). Libraqire Gallimard, Paris (1976).

7. S. Schütz-Bosbach, J. J. Musil, P. Haggard, Touchant-touché: The role of self-touch in the representation of body structure. Consciousness and cognition. 18, 2–11 (2009).

8. S. Gallagher, A. N. Meltzoff, The earliest sense of self and others: Merleau-Ponty and recent developmental studies. Philosophical psychology. 9, 211–233 (1996).

9. S. Dieguez, M. R. Mercier, N. Newby, O. Blanke, Feeling numbness for someone else’s finger. Current Biology. 19, R1108–R1109 (2009).

10. M. P. M. Kammers, F. de Vignemont, P. Haggard, Cooling the Thermal Grill Illusion through Self-Touch. Current Biology. 20, 1819–1822 (2010).

11. H. E. van Stralen, M. J. E. van Zandvoort, H. C. Dijkerman, The role of self-touch in somatosensory and body representation disorders after stroke. Philosophical Transactions of the Royal Society B: Biological Sciences. 366, 3142–3152 (2011).

12. M. Hara, P. Pozeg, G. Rognini, T. Higuchi, K. Fukuhara, A. Yamamoto, T. Higuchi, O. Blanke, R. Salomon, Voluntary self-touch increases body ownership. Frontiers in psychology. 6, 1509 (2015).

13. J. J. Gibson, The senses considered as perceptual systems. (1966).

14. R. Descartes, Treatise of man (Harvard University Press, 1972).

15. W. Penfield, T. Rasmussen, The cerebral cortex of man; a clinical study of localization of function. (1950).

16. B. Bridgeman, in The Oxford Handbook of Eye Movements (Oxford University Press, Oxford, 2011), pp. 511–521.

17. R. H. Lotze, Medicinische Psychologie oder Physiologie der Seele. Von Dr. Rudolph Hermann Lotze Professor in Göttingen (Weidmann’sche Buchhandlung, 1852).

18. W. M. Wundt, Beiträge zur Theorie der Sinneswahrnehmung (CF Winter, 1862).

19. N. Gangopadhyay, J. Kiverstein, Enactivism and the unity of perception and action. Topoi. 28, 63–73 (2009).

20. J. K. O’Regan, A. Noë, A sensorimotor account of vision and visual consciousness. Behavioral and brain sciences. 24, 939–973 (2001).

21. J. M. Findlay, I. D. Gilchrist, Active vision: The psychology of looking and seeing (Oxford University Press, 2003).

22. M. A. Goodale, A. D. Milner, others, Separate visual pathways for perception and action (1992).

23. H. C. Dijkerman, E. H. De Haan, Somatosensory processing subserving perception and action: Dissociations, interactions, and integration. Behavioral and brain sciences. 30, 224–230 (2007).

24. M. O. Ernst, M. S. Banks, Humans integrate visual and haptic information in a statistically optimal fashion. Nature. 415, 429–433 (2002).

25. H. Helson, S. M. King, The tau effect: an example of psychological relativity. Journal of Experimental Psychology. 14, 202–217 (1931).

26. W. J. Conover, Practical nonparametric statistics (John Wiley & Sons, 1998), vol. 350.

27. M. O. Ernst, H. H. Bülthoff, Merging the senses into a robust percept. Trends in cognitive sciences. 8, 162–169 (2004).

28. K. P. Körding, D. M. Wolpert, Bayesian integration in sensorimotor learning. Nature. 427, 244–247 (2004).

29. D. Alais, D. Burr, The Ventriloquist Effect Results from Near-Optimal Bimodal Integration. Current Biology. 14, 257–262 (2004).

30. H. H. Ehrsson, N. P. Holmes, R. E. Passingham, Touching a Rubber Hand: Feeling of Body Ownership Is Associated with Activity in Multisensory Brain Areas. J. Neurosci. 25, 10564–10573 (2005).

31. L. Dupin, P. Haggard, Dynamic Displacement Vector Interacts with Tactile Localization. Current Biology. 29, 492–498.e3 (2019).

32. S.-J. Blakemore, D. M. Wolpert, C. D. Frith, Central cancellation of self-produced tickle sensation. Nature neuroscience. 1, 635 (1998).

33. S. E. Laskin, W. A. Spencer, Cutaneous masking. I. Psychophysical observations on interactions of multipoint stimuli in man. Journal of Neurophysiology. 42, 1048–1060 (1979).

34. J. C. Craig, Attending to two fingers: Two hands are better than one. Perception & Psychophysics. 38, 496–511 (1985).

35. L. Tamè, C. Braun, N. P. Holmes, A. Farnè, F. Pavani, Bilateral representations of touch in the primary somatosensory cortex. Cognitive Neuropsychology. 33, 48–66 (2016).

36. E. B. de Condillac, Traité des sensations (chez les Librairies Associés, 1793).

37. A. Bicanski, N. Burgess, Neuronal vector coding in spatial cognition. Nature Reviews Neuroscience. 21, 453–470 (2020).

38. M. Martin, in The Contents of Experience, T. Crane, Ed. (New York: Cambridge University Press, 1992).

39. F. Fardo, B. Beck, T. Cheng, P. Haggard, A mechanism for spatial perception on human skin. Cognition. 178, 236–243 (2018).

40. W. James, The principles of psychology, Vol I. (Henry Holt and Co, New York, NY, US, 1890), The principles of psychology, Vol I.

41. P. Haggard, The Neurocognitive Bases of Human Volition. Annual Review of Psychology. 70, 9–28 (2019).

42. B. Hommel, R. W. Wiers, Towards a Unitary Approach to Human Action Control. Trends Cogn Sci. 21, 940–949 (2017).

43. M. Tsakiris, G. Prabhu, P. Haggard, Having a body versus moving your body: How agency structures body-ownership. Consciousness and Cognition. 15, 423–432 (2006).

44. P. Haggard, S. Clark, J. Kalogeras, Voluntary action and conscious awareness. Nat Neurosci. 5, 382–385 (2002).

45. R. Boehme, S. Hauser, G. J. Gerling, M. Heilig, H. Olausson, Distinction of self-produced touch and social touch at cortical and spinal cord levels. PNAS. 116, 2290–2299 (2019).

46. T. Metzinger, in Neural Correlates of Consciousness: Empirical and Conceptual Questions, T. Metzinger, Ed. (MIT Press, 2000; http://sammelpunkt.philo.at/267/).

47. V. Chambon, H. Théro, M. Vidal, H. Vandendriessche, P. Haggard, S. Palminteri, Information about action outcomes differentially affects learning from self-determined versus imposed choices. Nature Human Behaviour, 1–13 (2020).

48. J. Cohen, Statistical power analysis for the behavioral sciences 2nd edn (Erlbaum Associates, Hillsdale, 1988).

49. F. Faul, E. Erdfelder, A. Buchner, A.-G. Lang, Statistical power analyses using G* Power 3.1: Tests for correlation and regression analyses. Behavior research methods. 41, 1149–1160 (2009).

50. J. G. Eisenhauer, Regression through the origin. Teaching statistics. 25, 76–80 (2003).

51. A. Y. Leib, A. Kosovicheva, D. Whitney, Fast ensemble representations for abstract visual impressions. Nature communications. 7, 13186 (2016).

