## Supplementary Material for "Constructing spatial perception through self-touch"

Supplementary Materials

### **List of Supplementary Materials**

- Supplementary Methods: Estimated lag between the leader and follower robot trajectories
- Supplementary Results: Active conditions in Experiments 4-5
- Supplementary Figure S1: Correlation between weights (ω) in the two types of movement (active/passive) in Experiment 3
- Supplementary Figure S2: Correlation between weights (ω) in the two types of task (judge touch/movement) in Experiment 3
- Supplementary Table S1: ANOVA table of Experiment 1
- Supplementary Table S2: ANOVA table of Experiment 2
- Supplementary Table S3: ANOVA table of Experiment 3
- Supplementary Table S4: ANOVA table of Experiment 4
- Supplementary Table S5: ANOVA table of Experiment 5
- Supplementary Table S6: Normality data of Experiments 1-5
- Supplementary Video S1: Experimental setup, procedure, and example of three trials with different motor:tactile gains
- Supplementary Data File S1: Tables of means and individual weights (ω)

### **Supplementary** Methods: Estimated lag between the leader and follower robot trajectories

The Geomagic Touch X robotic arms we used provide movement monitoring within an event loop at a frequency of approximately 950-1100 Hz. However, to estimate the actual lag in our leader-follower system, we performed a control analysis on kinematic data from an experimental database with a similar design. We measured the time taken for the follower device to reach successively sampled positions along the forward movement axis of the leader device, in each trial of the experimental dataset. The mean lag was 2.47 ms (SD across 24 participants: 0.62 ms).

### **Supplementary** Results: Active conditions in Experiments 4-5

In the active conditions of Experiments 4 and 5 the weight of the interfering information was significantly different from 0 (*Z* < -3.061, *p* = 0.002, *r* < -0.883, in both cases; and 1 (*Z* < -3.061, *p* = 0.002, *r* < -0.883, in both cases; Bonferroni adjusted alpha for two multiple comparisons, against 0 and 1: α = 0.025 per test: Judge Touch – Active: median ω = 0.55 [95% Confidence Interval of the median = 0.46, 0.69]; Judge Movement – Active: 0.20 [0.14, 0.38]) (see Figure 4C), replicating the results from Experiments 1-3.

### **Supplementary** Figure S1: **Correlation between weights (ω) in the two types of movement (active/passive) in Experiment 3**

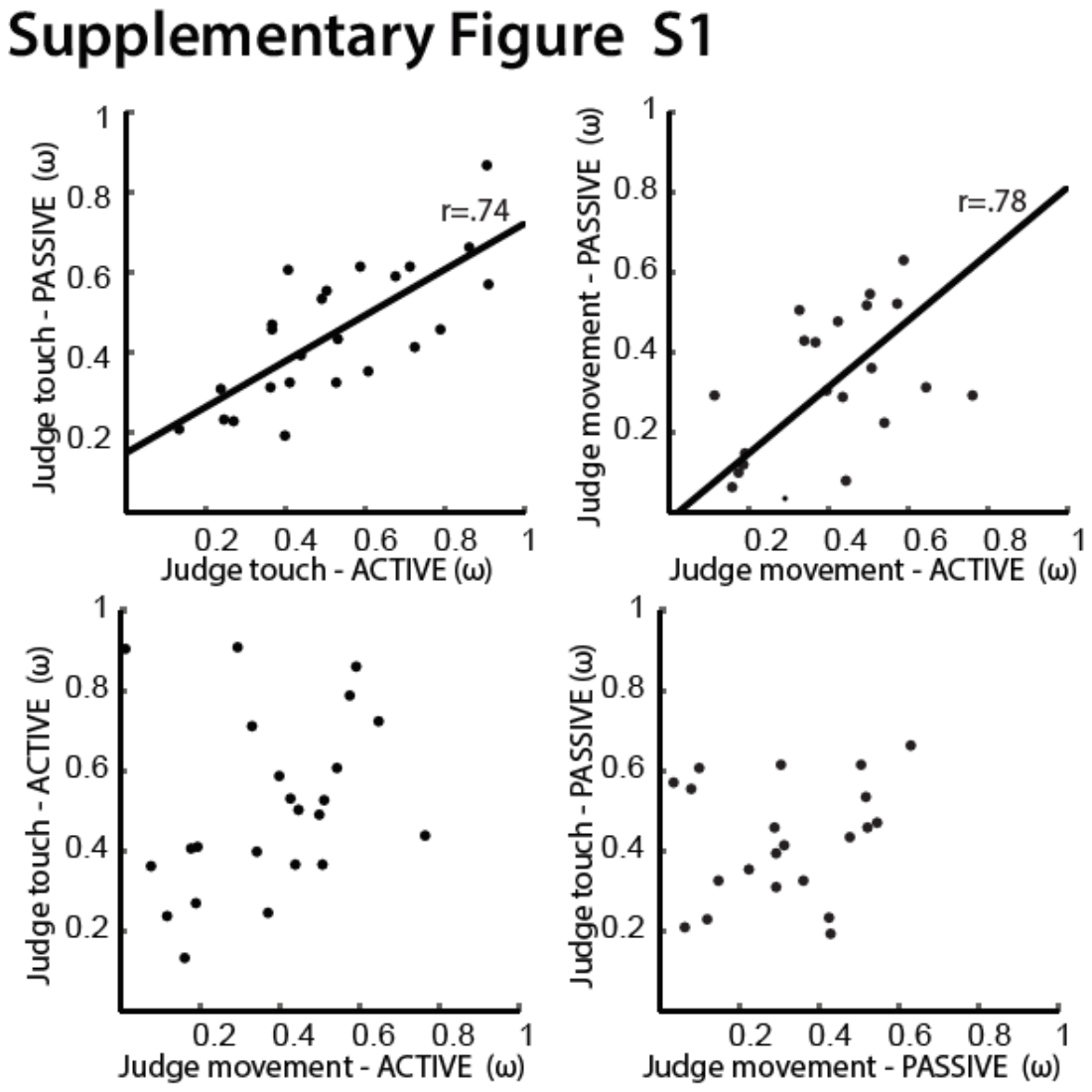

The within-subject design in Experiment 3 allowed us to investigate the correlation of motor and tactile weightings between all conditions (task type: judge touch/movement; movement type: active/passive). We reasoned that if the weights of task-irrelevant movement on touch judgements, and of task-irrelevant touch on movement judgements are correlated, it would mean that both reciprocal influences reflect a common cognitive process. For example, a positive correlation could indicate that people differ in the extent to which they express a general factor that determines perceptual selectivity vs perceptual interference between signals, without regard to the direction of interference. This factor might be called “selectivity of attention”. A negative correlation, instead, could mean that one particular signal, either movement or touch, dominates for each individual. When each individual makes judgements of their particular non-dominant signal, there would be strong interference from the dominant signal, leading to high weights. When they make judgements of the dominant signal, there would be minimal interference from the non-dominant signal, leading to low weights. A negative correlation might reflect a factor of intersensory dominance.

However, we found no significant correlation when comparing weights *ω* of judge touch and judge movement tasks for either Active (r=0.3, p=0.16; left panel) and Passive (r=0.1, p=0.66; right panel) movement conditions. This result rules out a strong overlap of processes related to the two tasks despite the reciprocal influence of movement over touch and vice versa. Thus, we did not find strong evidence for a role of trait selectivity of attention, or trait intersensory dominance in motor:tactile interactions.

### **Supplementary** Figure S2: **Correlation between weights (ω) in the two types of task (judge touch/movement) in Experiment 3**

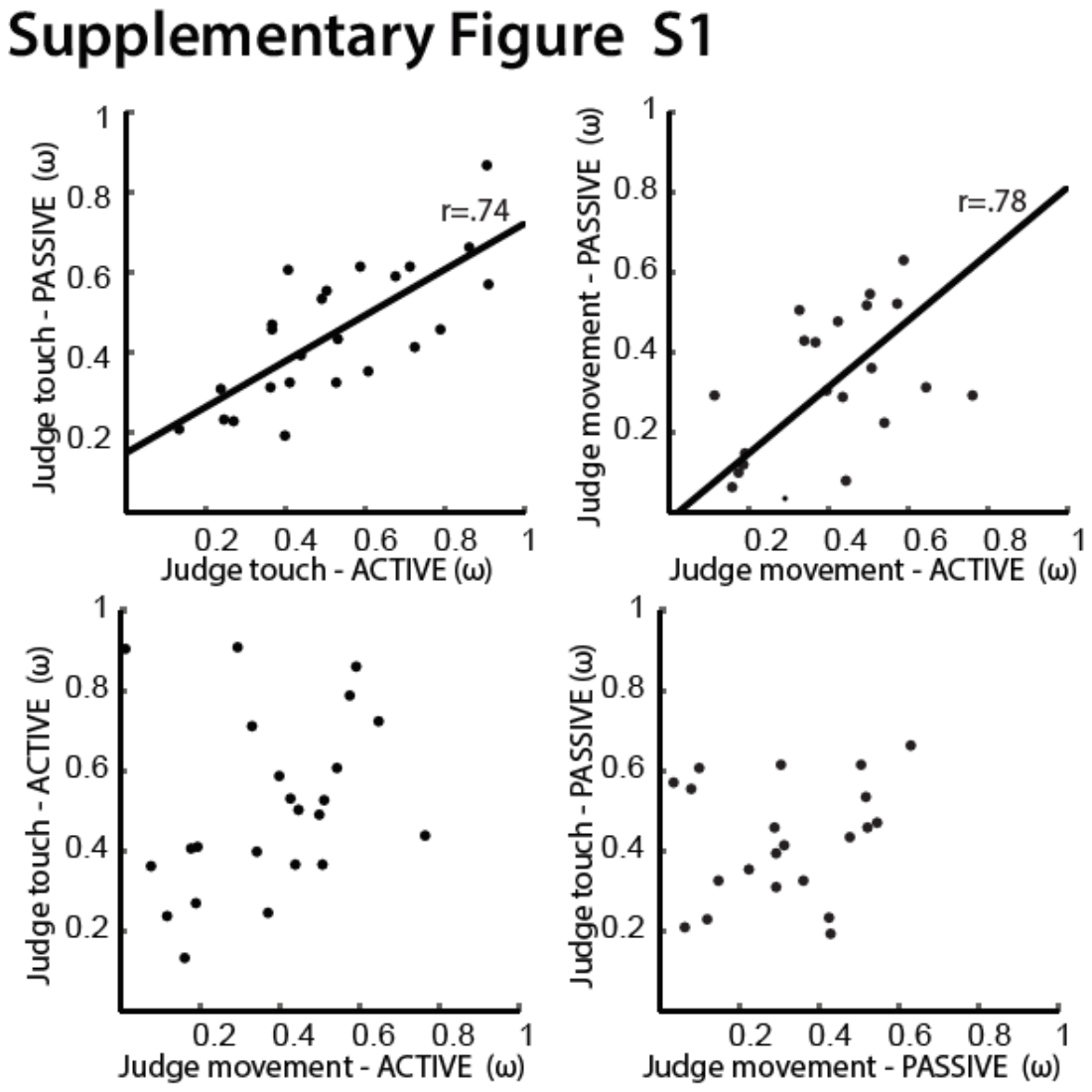

Next, we investigated the correlations between active and passive movement conditions in both tasks. We found a strong correlation of *ω* in Active and Passive movement conditions in judge touch (r = 0.74, p<0.001; left panel) and judge movement tasks (r = 0.78, p<0.001; right panel). This suggests that interference from movement when judging touch, and from touch when judging movement, could both involve some process that is common to both active and passive movements.

### **Supplementary Table S1: ANOVA table of Experiment 1**

|  | F | p | Eta |
| --- | --- | --- | --- |
| Mov. Type | .73 | .41 | .062 |
| **Extent** | **58.67** | **<.001** | **.84** |
| **Gain** | **28.71** | **<.001** | **.72** |
| Mov. Type x Extent | **4.4** | **.025** | **.29** |
| Mov. Type x Gain | 3.66 | .12 | .25 |
| **Extent x Gain** | **15.6** | **<.001** | **.59** |
| Mov. Type x Gain x Extent | .16 | .99 | .015 |

### **Supplementary Table S2: ANOVA table of Experiment 2**

|  | F | p | Eta |
| --- | --- | --- | --- |
| Mov. Type | .32 | .58 | .029 |
| **Extent** | **101.32** | **<.001** | **.90** |
| **Gain** | **34.54** | **<.001** | **.76** |
| Mov. Type x Extent | **21.52** | **<.001** | **.66** |
| Mov. Type x Gain | 4.73 | .003 | .30 |
| **Extent x Gain** | **6.80** | **.001** | **.38** |
| Mov. Type x Gain x **Extent** | .90 | .46 | .08 |

### **Supplementary Table S3: ANOVA table of Experiment 3**

|  | F | p | Eta |
| --- | --- | --- | --- |
| Task | .143 | .71 | .01 |
| Mov. Type | 1.61 | .22 | .07 |
| **Extent** | **219.1** | **<.001** | **.91** |
| **Gain** | **7.62** | **.007** | **.25** |
| Task x Mov. Type | .88 | .36 | .04 |
| **Task x Extent** | **7.92** | **.005** | **.26** |
| **Task x Gain** | **108.76** | **<.001** | **.83** |
| **Mov. Type x Extent** | **29.42** | **<.001** | **.56** |
| Mov. Type x Gain | .046 | .93 | .00 |
| **Extent x Gain** | **11.30** | **<.001** | **.33** |
| **Task x Mov. Type x Extent** | **4.82** | **.02** | **.17** |
| **Task x Mov. Type x Gain** | **5.75** | **.01** | **.20** |
| **Task x Gain x Extent** | **33.17** | **<.001** | **.59** |
| Mov. Type x Extent x Gain | 1.91 | .15 | .07 |
| Task x Mov. Type x Extent x Gain | .95 | .43 | .04 |

### **Supplementary Table S4: ANOVA table of Experiment 4**

The Condition factor has two levels: active or unimodal

|  | F | p | Eta |
| --- | --- | --- | --- |
| Mov. Type | **18.15** | **<.001** | **.62** |
| **Extent** | **729.6** | **<.001** | **.99** |
| **Gain** | **79.46** | **<.001** | **.88** |
| Condition x Extent | **12.49** | **.001** | **.53** |
| Condition x Gain | **104.75** | **<.001** | **.91** |
| **Extent x Gain** | **11.99** | **<.001** | **.52** |
| Mov. Type x Gain x Extent | **7.11** | **<.001** | **.39** |

### **Supplementary Table S5: ANOVA table of Experiment 5**

The Condition factor has two levels: active or unimodal

|  | F | p | Eta |
| --- | --- | --- | --- |
| Mov. Type | .38 | .55 | .03 |
| **Extent** | **81.99** | **<.001** | **.88** |
| **Gain** | **14.6** | **<.001** | **.57** |
| Mov. Type x Extent | .076 | .83 | .01 |
| **Mov. Type x Gain** | **9.98** | **<.001** | **.48** |
| Extent x Gain | 0.40 | .70 | .04 |
| Mov. Type x Gain x Extent | 1.41 | .26 | .11 |

### **Supplementary** Table S6: Normality data of Experiments 1-5

Kolmogorov-Smirnov normality tests (p-values)

| **Experiment** | **Condition** | **p-value** |
| --- | --- | --- |
| 1 | Active | **0.018*** |
| 1 | Passive | **0.013*** |
| 2 | Active | 0.65 |
| 2 | Passive | 0.15 |
| 3 | Judge Touch Active | 0.57 |
| 3 | Judge Touch Passive | 0.75 |
| 3 | Judge Movement Active | 0.67 |
| 3 | Judge Movement Passive | 0.84 |
| 4 | Active | 0.63 |
| 4 | Unimodal | 0.70 |
| 5 | Active | **0.04*** |
| 5 | Unimodal | 0.62 |

*The hypothesis of normality is rejected.

Since Experiment 1 and 2, and 4 and 5 are similar and that some conditions of experiment 1 and 5 are not normal, to be coherent, we used non-parametric tests (sign tests) for all the analyses of these experiments.

### **Supplementary** Video S1: Experimental setup, procedure, and example of three trials with different motor:tactile gains

A video depicting the main features of the experimental setup, the procedure, and an example of three trials with different motor:tactile gains can be found at: <https://tinyurl.com/yxf34yna>.

### **Supplementary Data File S1: Tables of means and individual weights (ω)**

The tables of means and individual weights for each experiment can be found at: <https://tinyurl.com/y3ssgho4>.
